# Single cell transcriptomic heterogeneity in invasive ductal and lobular breast cancer cells

**DOI:** 10.1101/2020.02.21.959023

**Authors:** Fangyuan Chen, Kai Ding, Nolan Priedigkeit, Ashuvinee Elangovan, Kevin M. Levine, Neil Carleton, Laura Savariau, Jennifer M. Atkinson, Steffi Oesterreich, Adrian V. Lee

## Abstract

Invasive lobular breast carcinoma (ILC), one of the major breast cancer histological subtypes, exhibits unique clinical and molecular features compared to the other well-studied ductal cancer subtype (IDC). The pathognomonic feature of ILC is loss of E-cadherin, mainly caused by inactivating mutations within the *CDH1* gene, but the extent of contribution of this genetic alteration to ILC-specific molecular characteristics remains largely understudied. To profile these features transcriptionally, we conducted single cell RNA sequencing on a panel of IDC and ILC cell lines, as well as an IDC cell line (T47D) with CRISPR-Cas9-mediated knock out (KO) of *CDH1*. Inspection of intra-cell line heterogeneity illustrated genetically and transcriptionally distinct subpopulations in multiple cell lines and highlighted rare populations of MCF7 cells highly expressing an apoptosis-related signature, positively correlated with a pre-adaptation signature to estrogen deprivation. Investigation of *CDH1* KO-induced alterations showed transcriptomic membranous systems remodeling, elevated resemblance to ILCs in regulon activation, and suggests *IRF1* as a potential mediator of reduced proliferation and increased cytokine-mediated immune-reactivity in ILCs.

## Introduction

Among subtyping systems of breast cancer, histological classification remains an essential criterion due to distinctive features of the major two subtypes—invasive lobular breast carcinoma (ILC) and invasive ductal breast carcinoma (IDC). ILC is the 6^th^ most common cancer in women, with an estimated 40,000 new cases in 2019, despite accounting for a smaller proportion of breast cancer cases (∼15%) compared to IDC (∼75%)^1^. ILC shows distinct signaling in pathways essential for breast cancer growth and proliferation compared to IDC – such as the WNT4 signaling in response to estrogen stimulus or blockade^2, 3^, increased PI3K/Akt signaling^4, 5^, enhanced IGF1-IGF1R activation^6^, and dependency on ROS1^7^, which suggest that ILC could benefit from unique treatment strategies. The most distinguishing molecular feature of ILC is loss of E-cadherin, largely arising from inactivating *CDH1* mutations. E-cadherin loss disrupts adherens junctions^8^ and leads to cells with a smaller and rounder morphology, a more scattered alignment within tumor stroma, and greater metastatic tropisms to ovaries, peritoneum or gastrointestinal (GI) tracts compared to IDC^9^. Such loss often couples with other molecular features, including the aberrant cytosolic localization of p120^10^. Meanwhile, E-cadherin-null tumor models also exhibit certain ILC resemblance: *in vivo*, the *TP53 CDH1* dual KO mouse model showed elevated anoikis resistance and angiogenesis as well as GI tract or peritoneum dissemination similarly to human cases^11^; while *in vitro*, hypersensitized PI3K/Akt signaling via GFR-dependent response was identified in both human and mouse ILC cell lines compared to their E-cadherin positive counterparts^4^. Despite numerous clinical observations and biological models, it is currently unclear how E-cadherin loss leads to many of the lobular-specific features.

Intra-tumor heterogeneity (ITH) is a hallmark of treatment resistance and mortality in cancer^12^. Multiple genetically distinct populations of cancer cells within the same tumor – typically arising from a series of mutational events—are dynamically selected by both intrinsic and external pressures and potentially preserve subclones with high invasiveness and/or drug resistance^13–16^. In addition to genetic diversity, transcriptional heterogeneity is also a major driver of ITH in multiple cancer types^17–20^. Such transcriptional variation, defined as cell states, appear transient and flexible in response to environmental stimuli while partially influenced by DNA alterations. Although ITH is frequently considered under *in-vivo* context, previous studies have shown there is considerable heterogeneity even for cell lines grown in culture. However, the extent of this intra-cell line heterogeneity in breast cancer models has not yet been comprehensively characterized^21, 22^.

To quantify ITH between cell lines, referred to as ICH (Inter-Cellular Heterogeneity), and investigate differences between IDC and ILC, we performed single cell RNA sequencing (scRNA-seq) on a panel of eight cell lines. We first investigated ICH in general: most cell lines consist of genetic subclones with unique copy number alterations (CNAs). Transcriptomic heterogeneity was shown for MCF7 cells specifically, revealing that it is dominated by cell cycle, in which cells dynamically transit through several well-defined phases. Despite the majority of cycling cells, a rare population exists distinctively outside the cell cycle with a non-transiting ‘dormant’ state. Characterization of such ‘outliers’ uncovered a unique apoptotic signature, which correlates with other functionally related signatures of dormancy in other cell lines and tumors.

We further inspected transcriptomic alterations induced by loss of E-cadherin, using CRISPR-mediated *CDH1* KO in a commonly used IDC line, T47D. Simple deletion of *CDH1* caused cells to cluster independently from wild type (WT) T47D cells in two-dimensional (2D) UMAP embedding. Given such distinctive and systemic differences are likely mediated by transcriptional factors (TFs)^23^, we deduced regulon activation states from scRNA-seq data, which illustrated elevated resemblance towards two out of the three ILCs in *CDH1* KO versus WT cells. Among the TFs identified, we found a regulon of IRF1 activated by *CDH1* KO, which also show higher expression in luminal A ILC tumors than IDCs. While the mechanism whereby loss of E-cadherin activates IRF1 is not known, IRF1 regulon activation conforms to the less-proliferative, and potentially more immune-enriched^24^ features of the lobular subtype.

## Results

### scRNA-seq of breast cancer cell lines

To investigate the effect of loss of E-cadherin in ILC, we generated T47D cells with *CDH1* KO using CRISPR-Cas9, and ensured its depletion at the protein level. scRNA-seq was performed on this T47D KO strain and its parental WT strain, along with seven additional groups of cells: the IDC cell line MCF7, MCF7 with *ESR1* Y537S mutation, referred to as MCF7-mut; three ILC cell lines: MDA-MB-134-VI, SUM44-PE, BCK4; as well as two immortalized but non-cancerous cell lines: the breast MCF10A, and human embryonic kidney 293 (HEK293) cells. Each cell line was cultured separately in standardized conditions, mixed at similar number for standard 10X chemistry v3 library preparation, and sequenced with NovaSeq 6000 system (Fig. 1a).

**Fig. 1.**
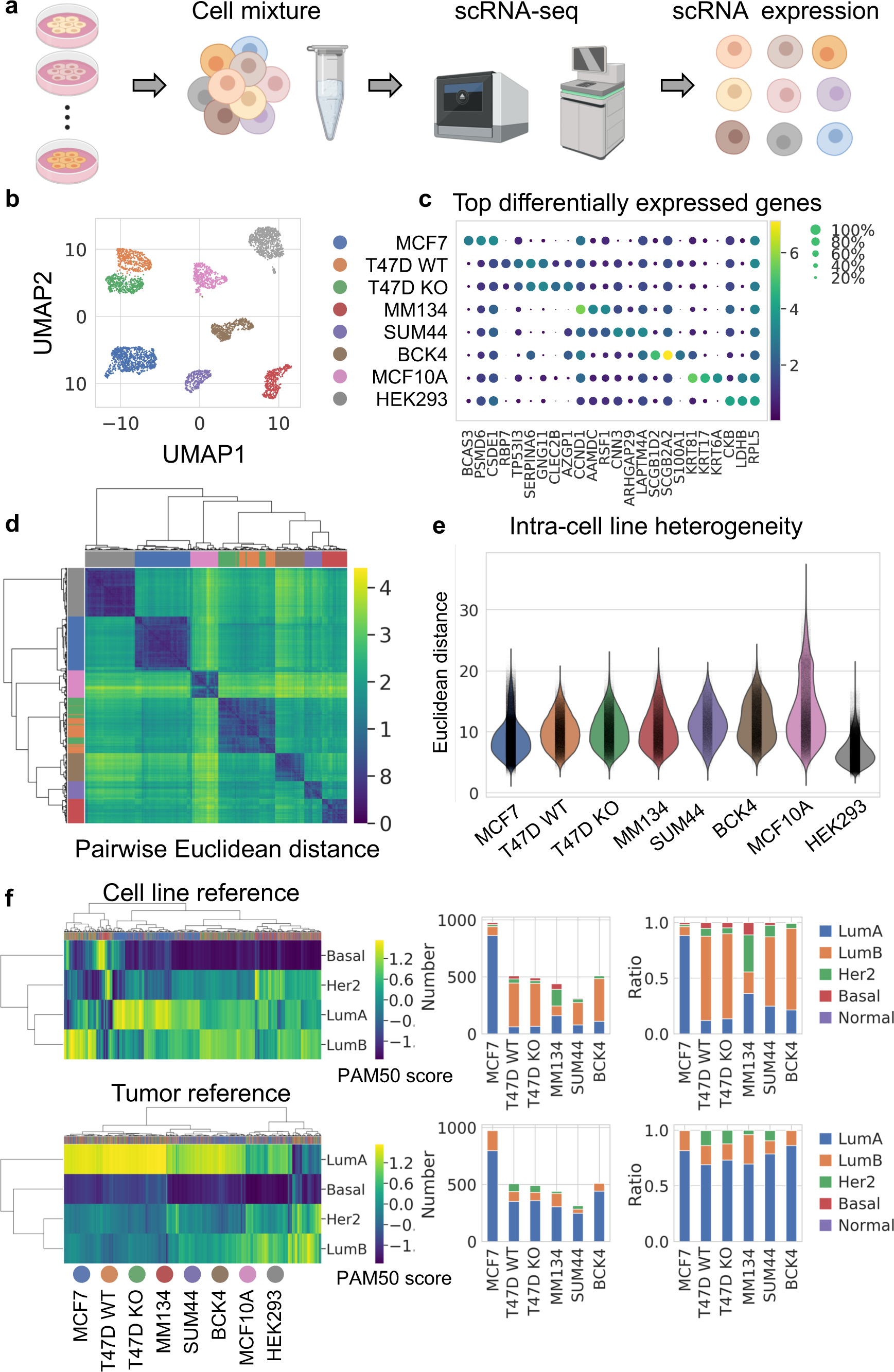
scRNA-seq of breast cancer and non-cancerous cell lines. a. Schematic pipeline of scRNA-seq. b. UMAP embeddings of 4,614 single cells in 8 clusters. Deconvoluted cell line identities are displayed in the same row as c. Number of cells: MCF7 (n=977), T47D WT (n=509), T47D KO (n=491), MM134 (n=439), SUM44 (n=314), BCK4 (n=512), MCF10A (n=491), HEK293 (n=881). c. Marker gene expression of each cell line. Top three differentially expressed genes were plotted for each cell cluster which had the smallest FDR when compared with all other clusters (Wilcoxon test, Benjamini-Hochberg (BH) adjustment). Every dot is colored by average expression of the gene and sized by the fraction of cells expressing the gene within that cell line. d. Hierarchical clustering (Euclidean distance, Ward’s method) of intercellular distances. X_i,j_, in the matrix represents the Euclidean distance between cell i and cell j using the top 30 principle components from the original expression matrix. Corresponding cell lines are colored on side bars, with the same color scheme as in b, c. e. Intercellular distances between every two single cells (calculated as Euclidean distance in d) within cell lines. f. Prediction Analysis of Microarray 50 (PAM50) subtypes scores (left) and assignment (right) of every single cell, using typical cell lines (upper) or estrogen-positive tumors (lower) as reference. Corresponding cell lines are colored on top bar of heatmap. Bar plots showed both absolute number of cells or the ratio of each PAM50 subtypes. LumA: luminal A, LumB: luminal B, Her2: HER2-enriched, Basal: basal-like, Normal: normal-like.

Dimensional reduction in 2D UMAP revealed eight distinct clusters (Fig. 1b). We deconvoluted single cell identities by mapping transcriptomes of each cluster to six cell lines with available bulk-RNA reference. Except for cluster 2, every cluster showed distinctive similarity to a specific bulk transcriptome, and thus identity was confidently assigned (Supplementary Figure 1). Cluster 2 was by default assigned as BCK4, which has no bulk RNA-seq data, and this identity was further confirmed by the exclusively high mucin expression in this cell line (Supplementary Figure 1). T47D *CDH1* KO and WT cells were two proximal but discrete clusters and showed altered E-cadherin expression as expected (Supplementary Figure 1). In contrast, the MCF7-mut and WT cells, despite being equally mixed, did not cluster separately, indicating limited transcriptomic differences when grown in standard media without estrogen deprivation. As these cells couldn’t be separated, we refer to them hereafter as MCF7 cells. After data pre-processing, the final single cell library consisted of 4,614 cells, approximately 500 cells for each cluster except MCF7 and HEK293, both of which contain approximately 900 cells (Fig. 1b).

Expression of key breast cancer genes were examined (Supplementary Figure 1), including hormone receptors (*ESR1* for estrogen receptor, *PGR* for progesterone receptor, *ERBB2* for human epidermal growth factor receptor 2), histology marker (*CDH1* for E-cadherin) and proliferation indicator (*MKI67* for Ki67). Consistent with the previous characterizations^25–27^, *ESR1* was expressed higher in the six breast cancer cell lines compared to MCF10A or HEK293; *PGR* showed high expression in T47D and BCK4 cells and low to medium in others; and *ERBB2* expression was higher in BCK4 than other cell lines. Consistent with histological classification, *CDH1* was highly expressed in IDC cell lines (MCF7, T47D WT) and MCF10A, compared to ILC cell lines or HEK293. All cell lines had abundant expression of *MKI67*. Despite cell-line specific expression, all markers showed a large variation in RNA abundance. Such heterogeneity is also reflected in PAM50 subtypes, calculated for each single cell with the subgroup-specific gene-centering method^28^ (Fig. 1f) – each cell line exhibits several PAM50 calls in spite of the luminal subtype dominance.

To quantify inter and intra-cell line differences, we calculated Euclidean distance among all pairs of single cells (Fig. 1d). Cells from each cell line showed greater similarity to each other, and T47D WT and KO were highly similar to each other. This analysis highlighted the intrinsic inter-cell line distinctions. Intra-cell line distances were also selected and compared, revealing intra-cell line heterogeneity which was highest in MCF10A, relatively consistently among breast cancer cells, and lowest in HEK293 (Fig. 1e).

### Inferred copy number aberrations (CNAs) reveal intra-cell line subpopulations that in part account for transcriptional heterogeneity

Cell lines are the most widely used laboratory model of cancer. However, studies have shown dissimilarities among breast cancer cell lines from different laboratories, potentially as a result of different culture conditions and/or intrinsic evolution of cells with genomic instability^21^. Even within a single cell line, transcriptomic subpopulations exist – composing a small or median proportion of the whole population, and are only partially explained by CNA^22^.

We examined genetic heterogeneity in cell lines using CNA inferred from scRNA-seq, a method described in multiple previous studies^18, 29^. To test robustness and accuracy of this method, we incorporated two external 10X scRNA-seq datasets, which investigated the same cell lines (MCF7 and T47D cells cultured in standard media^29^, plus three different MCF7 strains^21^). Different strains of the same cell type exhibited high resemblance to each other, as shown by co-clustering of T47D *CDH1* WT and KO cells with the external T47D dataset (Fig. 2a). Similarly, our MCF7 cells clustered with MCF7 strains from two other studies, in a different hierarchical branching to T47Ds. Three ILC cell lines (MDA-MB-134-VI, BCK4 and SUM44-PE) clustered together in a third independent branch from the hierarchical tree.

**Fig. 2.**
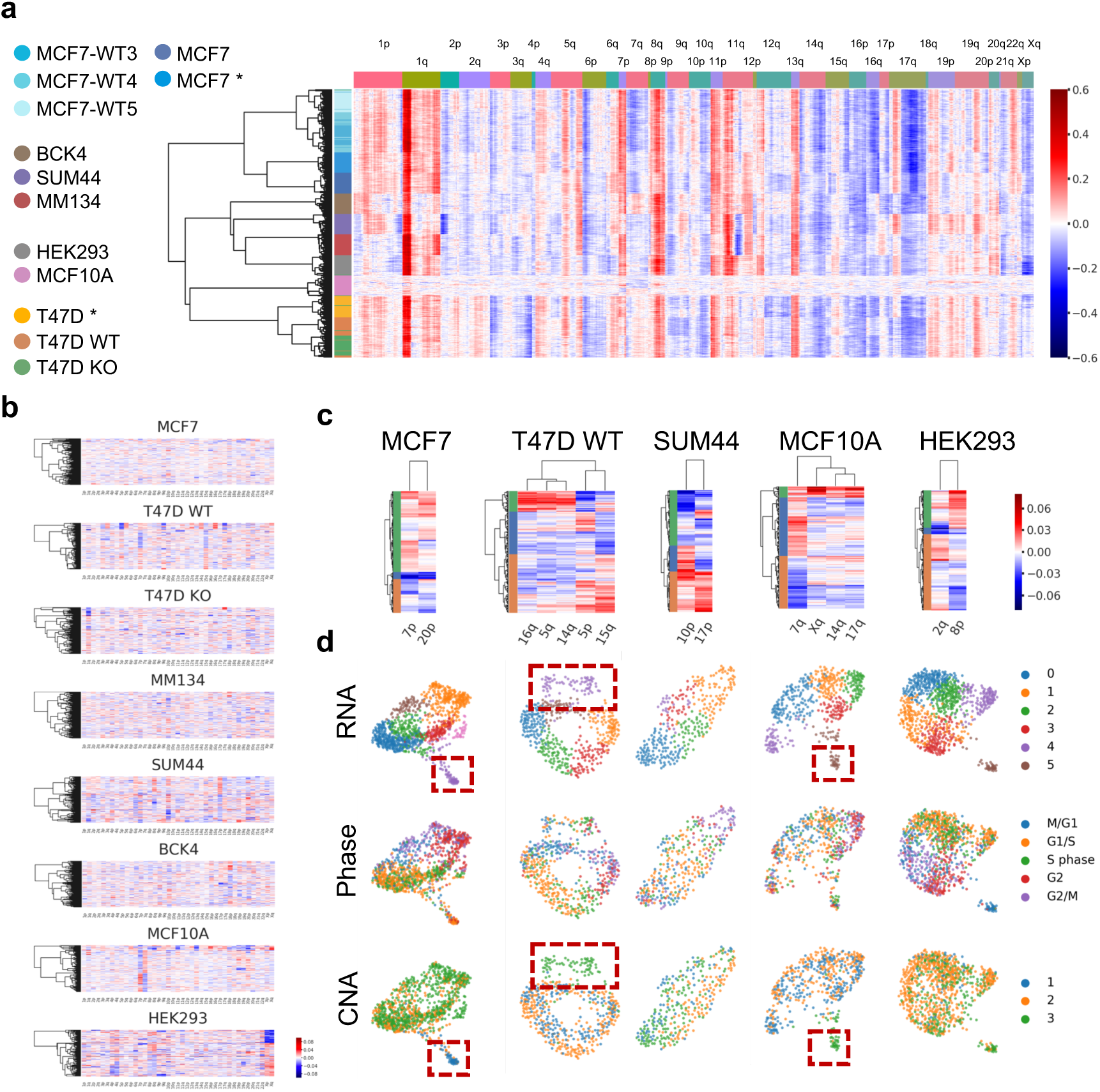
Intra-cell line subpopulations from inferred CNA. a. Copy number alteration (CNA) inferred from scRNA-seq of in-house and external cell lines (MCF7*, T47D*: MCF7 and T47D cells cultured in regular media^29^; MCF7-WT3/4/5: three MCF7 strains^21^), using average CNA of MCF10A as reference. 300 randomly selected cells for each cell strain were illustrated with hierarchical clustering (Euclidean distance, Ward’s method). b. Inferred CNA averaged at chromosome arm level. Only arms with more than 100 genes expression were selected. c. Cell lines with identifiable intra-cell line CNA subpopulations based on selected chromosome arms, colored on heatmap side bars. d. Intra-cell line RNA and CNA subpopulations, and cell cycle of cell lines in c. Clusters recurrently identified by both CNA and RNA are marked with squares.

To characterize genetic ICH, we identified subpopulations from CNA using selected chromosome arms as described by Kinker et al.^22^ (Fig. 2b,c; Supplementary Figure 2). For cell lines exhibiting CNA subclones, we compared the genetic subclones with transcriptomic subclones (derived from Louvain clustering from normalized RNA expression, and annotated by phases deduced from cell cycle gene expression) (Fig. 2d). Most of the cell lines investigated (5 out of 8) showed distinct genetic subpopulations not attributed to the cell cycle or scRNA-seq library quality (Fig. 2d, Supplementary Figure 2). Some but not all CNA clusters (CNA cluster 1 in MCF7, CNA cluster 3 in T47D WT and MCF10A), taking up a minority of the total population, corresponded to a transcriptomic subcluster based on both 2D layout and Louvain clustering (Fig. 2d).

### Apoptotic signature derived from a dormant-like MCF7 subpopulation

To investigate transcriptomic ICH, we focused on MCF7 cells, which had sufficient cell numbers in favor of statistical analysis. To cluster cells and select indicative features simultaneously, we used non-negative matrix factorization (NMF), which generated a matrix highlighting three major blocks of cells with corresponding genes, referred to as NMF clusters/genes (Fig. 3a, Supplementary Table 2).

**Fig. 3.**
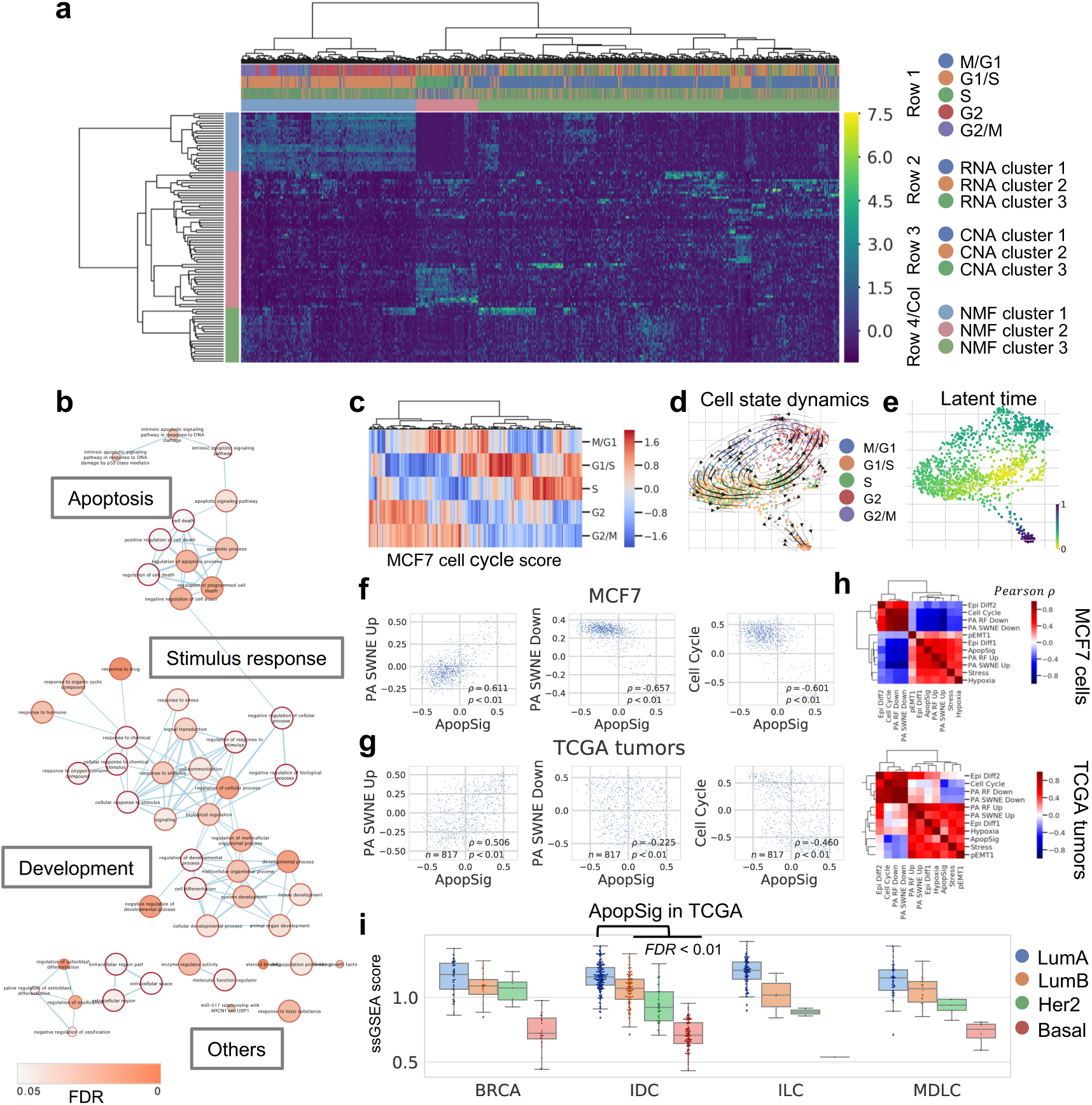
Transcriptomic heterogeneity in MCF7 cells. a. Clustering by Non-negative Matrix Factorization (NMF) in MCF7 cells (n=977). The first three rows of top bar showed respectively: cell cycle (row 1), RNA clusters (Louvain method, three clusters at resolution=0.4) (row 2) and CNA clusters (row 3). NMF clusters of cells and corresponding genes are shown in row 4 of top bar and the side bar. b. GO enrichment of marker genes of NMF cluster 2 cells (pink side bar in Fig. 3a). Terms connections based on similarity; nodes colored by enrichment FDR (over-representation test, BH adjustment) in Cytoscape 3.7.1. c. Cell cycle phase scores among single cells in MCF7. d. Dynamical changes of cell states through the cell cycle. Cells are colored by the assigned phase in the force-directed graph drawing 2D layout. Arrows show directions of cell state transition from RNA velocity analysis. e. Latent time among MCF7 cells from RNA velocity analysis, indicating developmental stages. f. Co-expression of GSVA scores of ApopSig with selected signatures in MCF7 cells (n=977) (PA SWNE Up: up-regulated gene signature in pre-adaptation^29^; PA SWNE Down: down-regulated gene signature in pre-adaptation^29^; Cell Cycle: cell-cycle related genes^18^). Correlation showed by Pearson *ρ* and p. g. Co-expression of GSVA scores of ApopSig with selected signatures (as in f) in TCGA breast tumors^32^ (n=817). Correlation showed by Pearson *ρ* and p. h. Pearson correlation of GSVA scores of ApopSig with selected signatures^18, 29^ in MCF7 cells (n=977) and primary breast tumors from TCGA (n=817). Hierarchical clustering was performed using Euclidean distance and Ward’s method. i. Single sample GSEA (ssGSEA)^48, 49^ scores of ApopSig in different subtypes of breast cancer from TCGA. (BRCA: breast cancer samples without histological annotation, IDC: invasive ductal carcinoma, ILC: invasive lobular carcinoma, MDLC: mixed ductal/lobular carcinoma). ApopSig ssGSEA scores are higher in LumA IDCs (n=200) than each of the other subtypes in IDC tumors (LumB: n=122, Her2: n=51, Basal: n=107) (FDR < 0.01, Wilcoxon test, BH adjustment).

Most cells belonged to NMF cluster 1 or 3, which overlapped with the two major RNA clusters (1 and 2 in Fig. 3a) or CNA subclusters (2 and 3 in Fig. 3a). A comparison with cell cycle showed that NMF cluster 1 correspond to the mitotic phase, while cluster 3 majorly consist of cells in G1/S, further supported by Gene Ontology (GO) enrichment (Fig. 3a, Supplementary Figure 3). The major effect of the cell cycle on transcriptional variation in MCF7 cells was demonstrated by highlighting the cell cycle phase of each cell (Fig. 3c), and the transition through different states predicted by RNA velocity analysis (Fig. 3d).

Despite the majority of cells apparently transiting through the cell cycle, there existed a minor population (NMF cluster 2) exhibiting a ‘dormant-like’ non-transiting state. This is consistently indicated by high latent time values from RNA velocity analysis (Fig. 3e). An inspection of highly expressed genes in this cluster revealed an enrichment of apoptosis-related pathways. We thus refer to this cluster of cells as Apop cells and their corresponding genes as ApopSig (signature). Interestingly, a recent report revealed that MCF7 cells contain a rare ‘pre-adapted endocrine resistant’ sub-population even when grown in regular media (DMEM, 10% fetal calf serum)^29^. A pre-adaptation signature (highly-expressed PA Up genes or lowly-expressed PA Down genes), derived from these cells, revealed a negative correlation with cell cycle and was indicative of dormancy. It is hypothesized that these pre-adapted cells may evade growth inhibition by anti-estrogens via exhibiting the less-aggressive dormant-like features. Motivated by this discovery, we investigated the association of the ApopSig with the pre-adapted signature. Despite a limited overlap in genes present in these two signatures (Supplementary Figure 3), ApopSig showed a significant correlation with both the PA Up (r=0.611, p<0.01) and PA Down signatures (r=-0.657, p<0.01) (Fig. 3f) in MCF7 cells. This correlation is similarly observed in TCGA breast tumors (Fig. 3g) or other breast cell lines (Supplementary Figure 3). Expression correlation with other functionally-relevant tumor signatures^18^ further illustrated a positive correlation of ApopSig with partial EMT (epithelial–mesenchymal transition), stress and hypoxia (Fig. 3h, Supplementary Figure 3). ApopSig showed enrichment in Luminal A tumors, which has the best prognosis among all PAM50 subtypes (Fig. 3i). High expression of ApopSig also indicates good prognosis, possibly due to its less aggressive manifestations, which holds true even when restricted to the estrogen receptor positive and LumA cohort (HR = 0.18, p = 0.022 by log rank test) (Supplementary Figure 3).

### *CDH1* point mutation results in loss of spliced E-cadherin RNA and alters expression programs related to the ILC phenotype

CRISPR-Cas9-mediated *CDH1* KO is induced by a single base pair deletion, which generated a premature stop codon, mimicking missense mutaitons found in ILC tumors and in genetically characterized ILC cell lines^30^. This point mutation led to depletion of both E-cadherin RNA and protein (Fig. 4a, Supplementary Figure 4), caused cells to lose cell-cell adhesion (Fig. 4a) and induced a profound effect on the transcriptome landscape, illustrated by distinct clustering of *CDH1* KO and WT cells (Supplementary Figure 4). When specifically examining spliced versus unspliced E-cadherin RNA abundance, the three ILC cell lines and T47D *CDH1* KO showed depleted spliced RNA abundance but comparable unspliced RNA distribution compared to MCF7 and T47D WT cells (Supplementary Figure 4). This suggests that *CDH1* mutation does not affect the nascent RNA transcript but post-transcriptional events, e.g., causing insufficient splicing or rapid degradation^31^. This result was mirrored in human tumors from TCGA^32^, in which IDC vs ILC showed a difference in exon RNA sequeuncing coverage than intron regions of *CDH1* (Supplementary Figure 4).

**Fig. 4.**
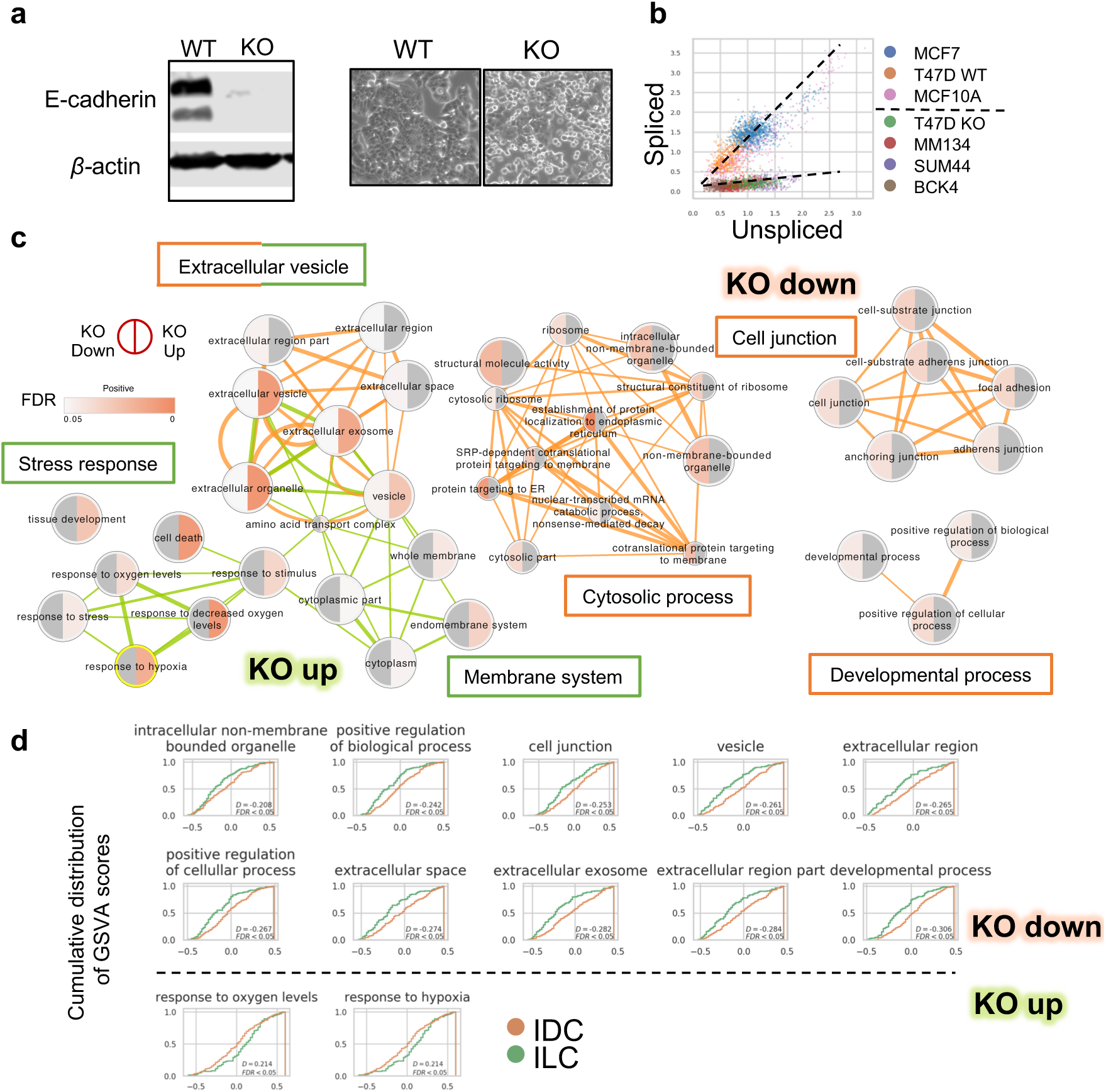
Differentially activated pathways in *CDH1* KO vs WT T47D cells and ILC vs IDC tumors. a. Left: western blot showing E-cadherin expression in T47D KO and WT cells. Right: morphology of WT and KO cells (10X bright field). b. Normalized unspliced and spliced *CDH1* RNA abundance among single cells. c. Enriched Gene Ontology terms of down (red linked) and up (green linked) regulated genes after *CDH1* KO in T47D cells. Terms connections based on similarity; nodes colored by enrichment FDR (over-representation test, BH adjustment) in Cytoscape 3.7.1. d. Cumulative distribution of GSVA scores of selected signatures in TCGA LumA IDC (n=200) and ILC (n=106) tumors. Right shifted curve indicates distribution of higher score values.

We explored what genes and pathways were differentially expressed (DE) after *CDH1* KO. As expected, cell junction-related components were down-regulated following loss of E-cadherin, along with some less specific pathways such as developmental or cytosolic processes (Fig. 4c). This is consistent with morphology changes in *CDH1* KO cells, which were more round with brighter margins, indicating decreased cell-cell contacts (Fig. 4a). Components in membranous system, endomembrane in particular, as well as stress response-related genes, were up-regulated in *CDH1* KO cells (Fig. 4c). Interestingly, extracellular vesicle-related pathways were enriched in both up and down regulated genes (Fig. 4c).

To investigate whether the transcriptomic changes in *CDH1* KO cells versus WT truly reflect ILC-IDC differences in tumors, we analyzed the expression of DE programs, refined by the overlap of DE genes with the original GO program (Supplementary Table 3), among the IDC and ILC in TCGA LumA cases. The majority of down-regulated gene sets (10 out of the 16 deduplicated gene sets) and some up-regulated ones (2 out of the 13 deduplicated gene sets) showed significant differences between ILC and IDC tumors, consistent with the trend in *CDH1* KO and WT models (Fig. 4d).

### An IRF1 regulon is activated following loss of E-cadherin, and is elevated in ILC

Loss of E-cadherin in epithelial cells has been reported to induce expression of multiple transcript factors (TFs) and trigger profound downstream phenotypic changes, such as metastasis promotion through epithelial-mesenchymal transition (EMT)^33^. While EMT does not seem to be a classical feature of ILCs^34, 35^, the vast transcriptomic changes in *CDH1* KO cells strongly suggest involvement of downstream TFs. We therefore searched for TF regulatory modules (regulons) which are increased or decreased in activity following *CDH1* deletion in T47D cells and investigated their expressions in ILC vs IDC tumors. Regulon activation profiles in each cell line was calculated using pySCENIC^36^. Commonly deduced regulons were binarized with an optimized threshold on AUC distribution and merged for all cell lines, which were used for hierarchical clustering (Fig. 5a,b, Supplementary Table 4). To more specifically quantify inter-cell line regulon activation differences, we measured the Jaccard Index between individual cells (Fig. 5c), where larger value indicates higher resemblance. Notably, T47D *CDH1* KO cells showed higher similarity to two out of three ILC cells (MDA-MB-134-VI, BCK4) than the two IDC cell lines (MCF7, T47D WT), (Fig. 5c, FDRs < 0.01 based on two sample K-S test, BH adjustment). This observation further supported that *CDH1* KO in IDC cells initiates an ILC specific TF regulon activation.

**Fig. 5.**
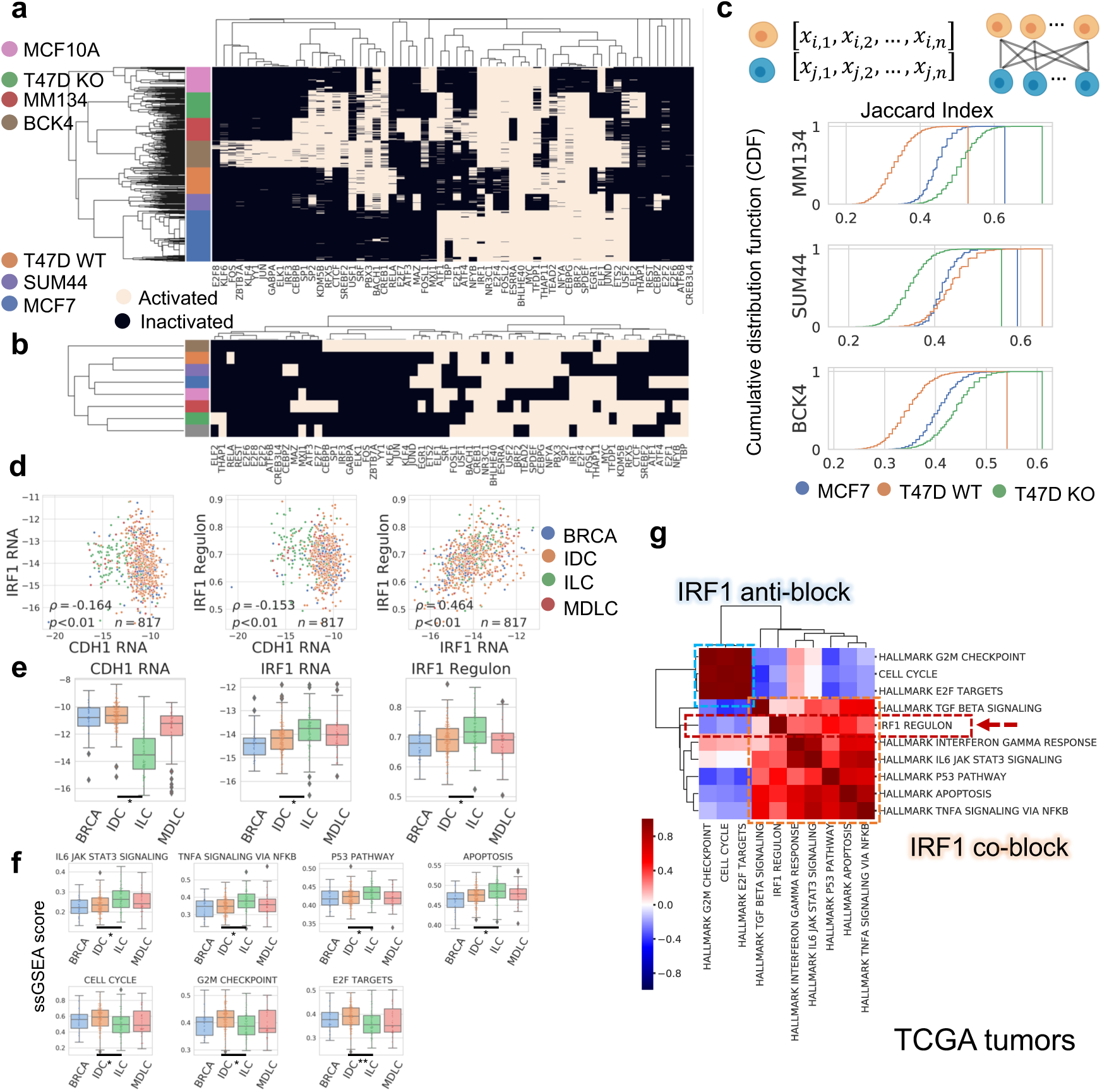
Regulon activation states in breast cell lines and TCGA tumors. a. Binarized regulon/TF activation profiles of each breast single cell deduced from scRNA-seq. b. Binarized regulon/TF activation profile for each breast cell line, based on the majority of single cell states in a. Hierarchical clustering by Jaccard distance, Ward method. c. Regulon activation similarity between each ILC cell line (reference) to MCF7, T47D WT and T47D KO (queries), quantified by Jaccard Index. For each reference cell line (per row, labeled on y axis), Jaccard Index was calculated between individuals in the reference population and every single cell of the three query breast cell lines respectively, depicted in cumulative distribution. Larger Jaccard Index indicates higher similarity. d. Co-expression patterns of *CDH1*, *IRF1* (log normalized) and IRF1 regulon (ssGSEA score) in TCGA LumA IDC and ILCs (*ρ*, Pearson correlation coefficient). e. Expression of *CDH1*, *IRF1* (log normalized RNA abundance) and IRF1 regulon (ssGSEA score) in TCGA LumA cases (BRCA: n=52, IDC: n=200, ILC: n=106, MDLC: n=57). Difference between IDCs and ILCs are significant (FDR<0.05) in all the three cases. f. ssGSEA scores of selected signatures in Fig. 5g which showed significant difference between TCGA LumA IDC (n=200) and ILC (n=106) tumors. g. Pearson correlation of ssGSEA scores of IRF1 regulon with relevant functional signatures in TCGA tumors (n=817). Signatures are divided to IRF1 co-block, which show positive correlation with IRF1 regulon; or IRF1 anti-block, which show negative correlation with IRF1 regulon.

We next identified regulons specifically activated following *CDH1* KO. Fourteen TFs were identified in this manner, which were further investigated regarding expression differences in LumA IDC and ILC in TCGA. Only IRF1 and CTCF showed significant differences (FDR<0.05), and only IRF1 exhibits higher expression in ILC (Supplementary Figure 5). Intriguingly, *IRF1* expression was also negatively correlated with *CDH1* in tumors (Fig. 5d), which further supports its activation in a lobular specific and E-cadherin associated manner. Similar observations were obtained in cell lines where ILCs generally have lower *CDH1* and higher *IRF1* or IRF1 regulon activation levels while IDCs show the opposite (except IRF1 regulon score of SUM44-PE, which is potentially due to influence of small sample size input to algorithm performance) (Supplementary Figure 5).

IRF1 is a canonical target of IFNγ, and in a pathway known to affect cell survival and proliferation. We next therefore examined co-expression of IRF1 regulon activation with selected MSigDB hallmark signatures with relevant functions in both tumors and cell lines (Fig. 5g, Supplementary Figure 5). Hierarchical clustering illustrated two distinct blocks, where IRF1 regulon positively correlates with IFNγ response, apoptosis, and signaling of TNF*α*, TGF*β* and IL-6; while showing a negative association with cell cycle (Fig. 5g, Supplementary Figure 5). Most pathways (4 out of 6) which were positively correlated showed enriched expression in ILC tumors given the difference is significant while all the three pathways with negative correlation showed the opposite (Fig. 5f).

## Discussion

scRNA-seq allows for single cell resolution of the transcriptome and is fundamentally altering our understanding of normal development and cancer. In this report, we used scRNA-seq to investigate inter-cellular heterogeneity of breast cancer cell lines, and specifically the unique features of ILC. scRNA-seq readily discerned differences between the cell lines, and genetic subclones were identified in most cell lines. Transcriptomic changes faithfully predicted the transition of cells through the cell cycle. However, in MCF7, a minor subpopulation of cells exist outside of the cell cycle, and these cells showed a dormancy related phenotype previously reported by other group^29^. ILC cell lines were distinct from IDC cell lines, and genetic deletion of *CDH1* caused transcriptional modeling in T47D as to be more similar to ILC than IDC cell lines. An investigation of activated regulons following loss of *CDH1* identified *IRF1*, which was also activated in LumA ILC.

scRNA-seq of cell lines revealed genetic and transcriptomic subpopulations within cell lines. A previous report of scRNA-seq in cell lines identified genetic and transcriptomic subpopulations in many cell lines, but not MCF7^22^. This inconsistency is unlikely due to strain artefacts, as our cell lines clustered correctly using CNA with the same cell lines from two other independent datasets, including the dataset which didn’t identify subclones in MCF7^22^. A possible reason is that we sequenced around five times the number of cells and thus had more power to find subpopulations. We found that MCF7 cells contained a subpopulation of non-cycling cells (Apop cells) with a dormancy phenotype reported by others^29^. Importantly, Apop cells corresponded to a subpopulation with pre-adaptation (PA) to endocrine therapy – also identified through scRNA-seq. The PA signatures are reported to support cancer survival in acute hormone deprivation. This strengthens the concept of transcriptionally-distinct minor subpopulations, which are present at all times, but in case of a harsh environment (e.g. hormone starvation), use their dormant phenotypes to survive and ultimately cause endocrine resistance.

scRNA-seq showed that IDC and ILC cell lines have distinct transcriptional programs, similar to tumors in TCGA; and that genetic loss of *CDH1* in an IDC cell line causes extensive transcriptional remodeling to make the resultant IDC *CDH1* KO cell line to resemble ILC, in both morphology and pathways. E-cadherin deficiency in lobular breast cancer was shown to be functionally associated with other structural proteins, e.g., elevated reliance on p120 in cytokinesis regulation^7^. From our data, we also observed structure-related transcriptomic changes after *CDH1* KO, such as junctional disruption; along with other features as expression increasement in endomembrane system, stress response and certain exocytosis pathways. These phenotypes from cell models were similarly identified when comparing clinical IDC and ILC LumA tumors.

The depletion of E-cadherin RNA and protein has been recognized in the majority of ILC tumors while promoter methylation is not associated with histological types^37^. This on one hand, justifies our use of cell lines for modeling ILC tumors, where MDA-MB-134-VI, SUM44-PE and T47D all harbor little methylation at *CDH1* promoter region (BCK4 had not been investigated)^38^; and on the other hand, suggests post-transcriptional modifications as potential driver of E-cadherin depletion. Our observation of alterations in *CDH1* spliced RNA, but not unspliced RNA in ILC from scRNA-seq data, provides evidence supporting this hypothesis. This was validated in TCGA bulk RNA-seq data via an approximation method of split exon/intron quantification, where we show more comparable intron RNA coverage in ILC as in IDC than exons. Notably, *CDH1* in T47D KO and ILCs all bear a pre-mature termination codon (PTC) while not necessarily contain disruptive mutations at splicing site (BCK4 mutation is currently unknown). In this context, loss of spliced mRNA is likely to result from the PTC-induced non-sense mediated decay, the main driver of E-cadherin transcript depletion as described in PTC-bearing gastric cancers^39^.

While E-cadherin is a membrane protein, its loss causes distinct transcriptional reprogramming, likely an indirect effect on TF activity, for example through inhibiting Kaiso’s TF activity as shown in mouse models^23^. To investigate this further, we examined regulon activation and identified an IRF1 regulon as being activated following *CDH1* KO, meanwhile showing higher RNA expression in ILC cell lines or tumors. As a tumor suppressor, IRF1 inhibits proliferation and prompts cell death. In breast cancer, IRF1 depletion could well indicate endocrine resistance, while its induction by IFN*γ* sensitize cancer cells to endocrine therapy^40^. These traits conform to multiple ILC phenotypes compared to IDC, e.g., being less proliferative and more apoptotic^41, 42^; and showing a better response to as well as a better outcome upon adjuvant endocrine therapy^43, 44^. Specifically, IRF1 mediates antiestrogen-induced apoptosis, by increasing expression of pro-apoptotic genes (BAK, BAX, BIK) while reducing that of anti-apoptotic genes (BCL2, BCLW, survivin)^45^. This corresponds to our observation of positive correlation of IRF1 regulon with hallmark apoptotic or p53 pathways, and the preferential activation of both pathways in ILC than IDC among LumA tumors. Apart from IFN*γ*, IRF1 can also be induced by other factors, such as IL-6, tumor necrosis factor (TNF) α and TGFβ^40, 46, 47^. Consistently, these pathways also correlate with IRF1 regulon through GSVA analysis (Fig. 5g, Supplementary Figure 5) while most of them showed enhanced signaling in ILCs (Fig. 5f), e.g., TNF*α* and IL-6 pathways. While pro-inflammatory signalings in tumor microenvironment has a complicated role in prognosis due to the pleiotropy of cytokines, they could reflect a coordination of enriched immune infiltration and/or enhanced immune reactivity in ILC tumors. Such immune signature enrichment, as has been shown previouslys^24^, might be predisposed by the E-cadherin mediated IRF1 activation within tumor cells and may suggest immune-sensitizing therapies in lobular breast cancer treatment.

In summary, scRNA-seq of breast cancer cell lines has revealed significant intra-cell line genetic and transcriptomic heterogeneity, with identification of dormant cells likely primed for anti-estrogen resistance. Knockout of *CDH1* in IDC mimics features of ILC and highlights the power of single cell sequencing to reveal unique features of breast cancer.

## Methods

### Generation of T47D *CDH1* KO cells

Knockout of *CDH1* was performed using CRISPR-Cas9 with the Gene Knockout Kit (V1) from Synthego (Redwood City, California). Four potential sgRNAs (1.CCGGTGTCCCTGGGCGGAGT, 2.CCTCTCTCCAGGTGGCGCCG, 3.GGCGTCAAAGCCAGGGTGGC, 4.CTCTTGGCTCTGCCAGGAGC) were selected based on sequence screening to target exons or introns of *CDH1* and introduce a protein truncating indel. Each sgRNA was introduced as an oligonucleotide with Cas9 2NLS Nuclease using nucleofection. Following a brief incubation period of each sgRNA with the Cas9, the ribonucleoprotein complex was nucleofected into T47D cells using the Lonza 4D-Nucleofector. 72 hours post nucleofection, half of the cell population was subjected to PCR for *CDH1* (F: 5’AGGAGACTGAAAGGGAACGGTG and R: 5’GTGCCCTCAACCTCCTCTTCTT) and sanger sequencing was used to confirm the presence of an indel. sgRNA 2 population turned out to induce the most complete protein depletion than other pools, and was thus chosen based on the sequencing results; demonstrating a 1bp deletion at exon 2 (c.321delC, in NM_004360.5), which caused a frameshift with a pre-mature stop codon at exon 16. To select pure KO clones, the sgRNA 2 cell population was single cell sorted into 96 wells by FACS, and supplemented with filtered T47D conditioned media. Upon colony formation, clones were expanded, and knockout success was examined by Sanger sequencing and Western blot to confirm protein loss (anti-*CDH1* antibody, BD #610182). 8 clones with the least E-cadherin protein expression by immunoblot were then pooled in equal ratio and named T47D *CDH1* KO. Images of KO and parental WT cells were obtained with 10X bright field using Olympus IX83 Inverted Microscope.

### Cell line preparation

MCF7, T47D, MDA-MB-134-VI, SUM44-PE, MCF10A and HEK293 were all purchased from American Type Culture Collection (ATCC) and identity authenticated by DNA fingerprinting (University of Arizona). Cells were routinely tested for mycoplasma and were negative at all times. MCF7 with *ESR1*-Y537S were generated previously^50^. BCK4 was a gift from Brita Jacobson (University of Colorado). Cells were maintained in media described in Supplementary Table 1.

### Single-cell RNA sequencing

Nine groups of cells, each with viability > 90% based upon Trypan Blue staining and Invitrogen automated cell counting, were fixed separately at equal number (round 1,000,000 cells per group) in 90% methanol at 4°C for 15 minutes and temporarily stored at −80°C. The cell suspension was rehydrated, mixed and processed following 10X Genomics 3’ Chromium v3.0 protocol at University of Pittsburgh Genomics Core. The library was sequenced with NovaSeq 6000 S1 flow cell at the UPMC Genome Center, getting around 400 million paired reads in total.

### scRNA-seq data pre-processing

Raw FASTQ data was aligned and quantified using GRCh38 reference with Cell Ranger (v3.0.2) (https://support.10xgenomics.com/single-cell-geneexpression/software/pipelines/latest/using/count) and velocyto CLI (v0.17.17)^51^. The resulting loom files were loaded with scVelo (v0.1.25)^52^ and processed with Scanpy (v1.4.4)^53^. Doublet removal was performed using Srublet^54^. Low quality cells and genes were filtered out by selecting cells expressing more than 2000 genes, having UMIs between 8000 and 10,000 with a mitochondrial gene percentage of less than 15%; and selecting genes with detectable expression in at least 2 cells, resulting in a final library of 4,614 cells and 21,888 genes. Quality metrics of the single cell library (number of genes, number of UMIs, and mitochondria reads percent for each cell) are depicted in Supplementary Figure 1.

From the filtered matrix, spliced and unspliced reads were normalized, converted to log scale, and imputed respectively by scVelo (v0.1.25)^52^, using scvelo.pp.normalize_per_cell, scvelo.pp.log1p and scvelo.pp.moments. The top 30 principle components were calculated using the 3000 most variable genes. This was followed by dimensional reduction using UMAP (scanpy.tl.umap) and clustering with Louvain method (scanpy.tl.louvain, resolution=0.2), which demonstrated eight distinct clusters in 2D UMAP embedding.

### Cell line identification

Bulk RNA-seq data were acquired from public database (MCF7, T47D, MDA-MB-134-VI from CCLE^55^; SUM44-PE^56^; MCF10A^57^; HEK293^58^). No bulk RNA-seq data of BCK4 was available. For the five breast cell lines, FASTQ files were obtained from Sequence Read Archive (SRP186687, SRP026537, SRP064259), quantified using Salmon (v0.12.0)^59^ using GRCh38 reference. For HEK293, RNA counts from GSM1867011 and GSM1867012 were directly downloaded and the average expression for all genes of these two sample were used subsequently. 1,444 genes were present in all the three data sources (breast cell lines, HEK293 as reference cell lines; and scRNA-seq (3,000 genes by 4,614 cells) as query cells), which were selected for further analysis.

Pearson correlation was calculated between each single cell in the 8 clusters and each reference cell line using log normalized counts. The reference with the most significantly right-skewed coefficient distribution was assigned to the query cluster (Supplementary Figure 1). Every cluster was successfully assigned except cluster 2, which was by default BCK4. This was further confirmed by the high levels of MUC2 expression (Supplementary Figure 1). WT and KO T47D cells were distinguished by *CDH1* expression (Supplementary Figure 1).

### PAM50 assignment

The six breast cancer cell lines (cell lines except MCF10A and HEK293) were classified with PAM50 subtypes with subgroup-specific gene-centering method^28^, using either tumor or cell line as reference. Normalized expression of genes in scRNA-seq data that overlapped with the PAM50 panel were selected and centered with the pre-calculated ER+ group-specific quantiles, as described by Zhao et al.^28^ For each single cell, a Spearman’s rank correlation coefficient was calculated between the centered expression vector and four PAM50 subtype (LumA, LumB, HER2, Basal) centroids, using either tumor data from the original University of Northern Carolina dataset, or bulk RNA-seq of representative cell lines generated as described above (LumA: MCF7, LumB: BT-474, HER2: SK-BR-3; Basal: MDA-MB-231, selected according to Roden et al.^60^). These correlation coefficients were standardized in each cell using z-score, and the subtype with highest correlation coefficient was assigned to this cell, which showed LumA/LumB dominance (Fig. 1f).

### CNA Inference

Two external datasets with scRNA-seq data of cell lines were integrated with our in-house data for CNA inference - MCF7 and T47D cells from Hong et al.^29^, and three MCF7 strains (WT-3,4,5) from Ben-David et al.^21^ A raw count matrix from Hong et al.^29^ was directly downloaded from GSE122743 while FASTQ files from Ben-David et al.^21^ were reprocessed to derive the count matrix, following steps described in the scRNA-seq data pre-processing section. The average expression vector of the in-house MCF10A cells was utilized as reference.

Expression count matrix of each cell strain was first log normalized, sorted by genes’ chromosomal coordinates, and then merged with others using commonly detected genes. 300 cells were randomly selected from each strain. Only chromosome arms containing more than 100 genes were kept, and single cell expression was averaged with a moving 100 gene window. This averaged expression was used to fit a linear regression model against the MCF10A reference vector, and the residual value was assigned as the inferred CNA (Fig. 2a).

### Identify cell subclusters from inferred CNA

To identify intra-cell line CNA subclusters we adopted a similar method as reported by Kinker et al.^22^ For each cell line, the log normalized expression matrix was centered for each gene and moving average of 100 gene window was calculated along the genome. For each chromosome arm, a gaussian mixture model (GMM, implemented in scikit-learn^61^) was fitted to the distribution. Selected chromosome arms should best fit with more than one component in GMM, each of which having more than 20 cells with confidence higher than 95%. These selected chromosome arms were then used for hierarchical clustering to identify subcluster of cells (Euclidean distance, Ward method) (Fig. 2c). The top three hierarchical branching were extracted and differentially colored as the potential CNA subpopulation.

### Identify cell subclusters from RNA

To identify intra-cell line transcriptomic subpopulations, RNA counts of each cell line were extracted from raw data, filtered, log normalized and imputed, using the top 5000 variable genes following steps described in the scRNA-seq data pre-processing section. Cells were laid out in a force-directed graph drawing implemented in Scanpy^53^. Louvain clustering was conducted in each cell line with resolution=1 in Fig. 2d as an empirical value; and with resolution = 0.4 for MCF7 in Fig. 3a, as to generate 3 clusters in correspondence to the 3 clusters via other classification methods.

### Cell cycle scoring

Five lists of phase marker genes (M/G1, G1/S, S, G2, G2/M) of the cell cycle were obtained^62^. Log normalized expression of each cell line was acquired as described in the last section. For each cell line and phase, correlation coefficients were calculated between each gene and the average gene set expression. Genes with the highest (the upper 40% quantile) coefficients were selected as the refined cell cycle markers for that cell line. A score for each phase was calculated for each cell, as the average expression of cell cycle markers minus that of a randomly selected gene set of the same size (implemented in Scanpy^53^), which is standardized firstly in each cell then in each phase by z-score (Fig. 3c). The phase with the highest score was assigned to the cell.

### RNA velocity analysis

For each cell line, the union of the refined cell cycle markers were selected to illustrate cell state transitioning dynamics, following the RNA velocity pipeline implemented in scVelo^52^ using stochastic mode, with latent time calculated under the same setting. Stream arrows indicating transition dynamics were depicted on the force-directed graph drawing of individual cell line.

### NMF clustering

NMF was performed as described by Puram, et al.^18^ in MCF7 cells. The log normalized expression matrix of 5000 genes and 977 cells was centered for each gene to obtain the relative expression (*E_i,j_*) for every gene i in each single cell j. Negative values was replaced by 0 and NMF was then conducted with k ranging from 5 to 15 (implemented in sklearn.decomposition.NMF). For each k, top 50 genes with highest D score^63^ were defined as a gene set. Recurrent gene sets, which have Jaccard Index > 0.6 with at least one another gene set, were selected and merged as indicative features (101 genes). *E_i,j_* of these 101 genes among MCF7 cells were standardized for each cell by cell-wise z-score and showed in heatmap, clustered for both genes and cells (Euclidean distance, Ward method) (Fig. 3a). Clades from the top three hierarchical branching were selected as NMF clusters.

### Differentially expressed genes

Differentially expressed genes (DEGs) for each cell line were derived by comparing this cell line versus all other cells using Wilcoxon test (BH adjustment, implemented in Scanpy^53^) (Fig. 1c). To calculate DEGs in T47D *CDH1* KO versus WT cells, the two groups were extracted from raw data, filtered, log normalized and imputed, using the top 3000 variable genes following steps described in the scRNA-seq data pre-processing section. DEGs were calculated comparing between each other (Wilcoxon test, BH adjustment). The top 100 genes with the smallest FDR were selected as marker genes for each group.

### Gene set enrichment analysis

Gene lists of interest (marker genes of MCF7 NMF clusters in Fig. 3; T47D WT and KO DEGs in Fig. 4) were submitted to the gProfiler website^64^ (https://biit.cs.ut.ee/gprofiler/) for pathway enrichment analysis using over representation test with default parameters. Enriched terms of GO dataset (Biological Process and Cellular Component) were selected as input in Cytoscape (v3.7.1)^65^ Enrichment Map, in GeneMANIA Force Directed Layout (similarity_coefficient mode) colored by FDR.

### Public gene sets

Gene sets used for GSVA scoring were obtained as follows: ApopSig derives from genes corresponding to NMF cluster 2 (Fig. 3a); PA SWNE Up or Down signatures, representing pre-adaption features, were accessed from Hong et al.^29^; hallmark gene sets in Fig. 5g were from MSigDB hallmark dataset^66^; and other intrinsic tumor signatures in Fig. 3h were from Puram et al.^18^ Refined GO terms in Fig. 4d are the overlap of the original GO term with DEGs between T47D WT and KO cells. Target genes in IRF1 regulon from ChEA or ENCODE datasets were downloaded from Enrichr libraries^67, 68^ (ChEA_2016, ENCODE_TF_ChIP-seq_2015).

### GSVA scoring in TCGA and METABRIC

Expression count matrix of TCGA breast cancer dataset was obtained from cBioportal^69, 70^ (meta_RNA_Seq_v2_expression_median.txt) and log normalized as 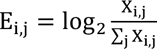 for sample i and gene j; corresponding breast cancer subtypes (PAM50, histology) were obtained from Ciriello et al.^37^ METABRIC^71^ RNA microarray data was downloaded from Synapse and log normalized. Gene set variation analysis (GSVA) of selected gene set was performed with GSVA R package^48, 49^, in ssgsea or gsva mode with default parameters. Comparisons between TCGA ILC versus IDC cases are all limited to LumA population unless otherwise specified.

For survival analysis, only estrogen receptor positive (ER+) and LumA patients were selected to avoid influence caused alone by PAM50 subtype. ApopSig ssGSEA scores were stratified into low and high levels with optimized threshold using in surv_cutpoint and surv_categorize implemented in R survminer package (https://cran.r-project.org/web/packages/survminer/index.html). Kaplan-Meier curve were plotted for each group using disease free survival, with p value from log-rank test, and hazard ratio (HR) calculated using univariate Cox regression (coxph in R survival package, https://cran.r-project.org/web/packages/survival/index.html).

### *CDH1* exon and intron coverage

TCGA RNA BAM files were accessed from The Pittsburgh Genome Resource Repository (https://www.pgrr.pitt.edu/). Bulk RNA-seq BAM files of breast cancer cell lines were generated as described in Tasdemir et al.^72^ RNA counts of each exon and intron of *CDH1* was quantified with bedtools (v2.29.1) counts mode and normalized by dividing the total number of counts within the sample (Supplementary Figure 4). One representative ILC and IDC patient from TCGA^37^, along with one cell sample from each cell line, were selected for visualization using Integrative Genome Viewer^73^, showing *CDH1* coverage from intron 11 to exon16 with GRCh38 as reference genome (Supplementary Figure 4).

### Regulon activation profiles

Regulon activities were inferred from raw expression counts of each cell line following the default pySCENIC^36^ pipeline (https://github.com/aertslab/pySCENIC). Each cell was eventually assigned with an AUC score for every regulon, indicating its activation status. These scores were then binarized into either “on” or “off” states, by setting an optimized threshold on the distribution of each regulon among all cells using the skimage.filters.threshold_minimum function. Target genes of each regulon were selected if it is commonly deduced under that TF in at least two of the following cell lines: T47D KO, MDA-MB-134-VI, SUM44-PE and BCK4 (Supplementary Table 4).

### Jaccard index

For every pair of cells, a Jaccard index was calculated using the two corresponding binarized regulon activation vector of the two cells, X and Y, as 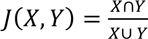, a large value of which indicates higher similarity. For every two cell lines, the Jaccard index of all pairwise combinations between the two population was selected and plotted as the cumulative distribution (Fig. 5c).

### Statistical analyses

All the analyses and plots were generated in Python (v3.7) (http://www.python.org) or R (v3.6) (www.r-project.org). All statistical tests are two-sided, unless specified otherwise.

### Code availability

Codes and important intermediate data will be available on github (https://github.com/leeoesterreich?tab=repositories) upon publication.

### Data availability

Processed files are deposited in Gene Expression Omnibus (GSE144320) and raw FASTQ files are deposited at SRP245420, which will be available upon publication. Processed data can also be accessed interactively through our server, implemented by SCelVis^74^, at http://167.172.151.214:8050/dash/viz/CellLines.

## Acknowledgements

This project benefited from resources provided by the University of Pittsburgh HSCRF Genomics Research Core, the UPMC Genome Center, the Pittsburgh Genomic Research Repository (dbGAP), and the University of Pittsburgh Center for Research Computing. The authors would like to thank Fangping Mu and Uma Chandran for bioinformatics assistance.

The China Scholarship Council and Tsinghua University provided financial support for F.C. This work was supported by the Breast Cancer Research Foundation [A.V.L. and S.O.]; Susan G. Komen Scholar awards [SAC110021 to A.V.L., SAC160073 to S.O.]; the National Cancer Institute [5F30CA203154 to K.M.L., P30CA047904]; Magee-Women’s Research Institute and Foundation, and the Shear Family Foundation. S.O. and A.V.L. are Hillman Fellows. The content is solely the responsibility of the authors and does not necessarily represent the official views of the National Institutes of Health or other Insitutes and Foundations.

## Author contributions

Study concept and design (NP, SO, AVL); acquisition, analysis, or interpretation of data (FC, KD, AE, NP, SO, AVL); drafting of the manuscript (FC, NP, SO, AVL); critical revision of the manuscript for important intellectual content (all authors); administrative, technical, or material support (AB, RJW, NT, DV, WH, JKK, NED, AMB, RS, MJS, ES, RJH, LY, CD, AM, BIH, JB, PCL).

