## Supplementary Figures for "Single cell transcriptomic heterogeneity in invasive ductal and lobular breast cancer cells"

### Slide 1
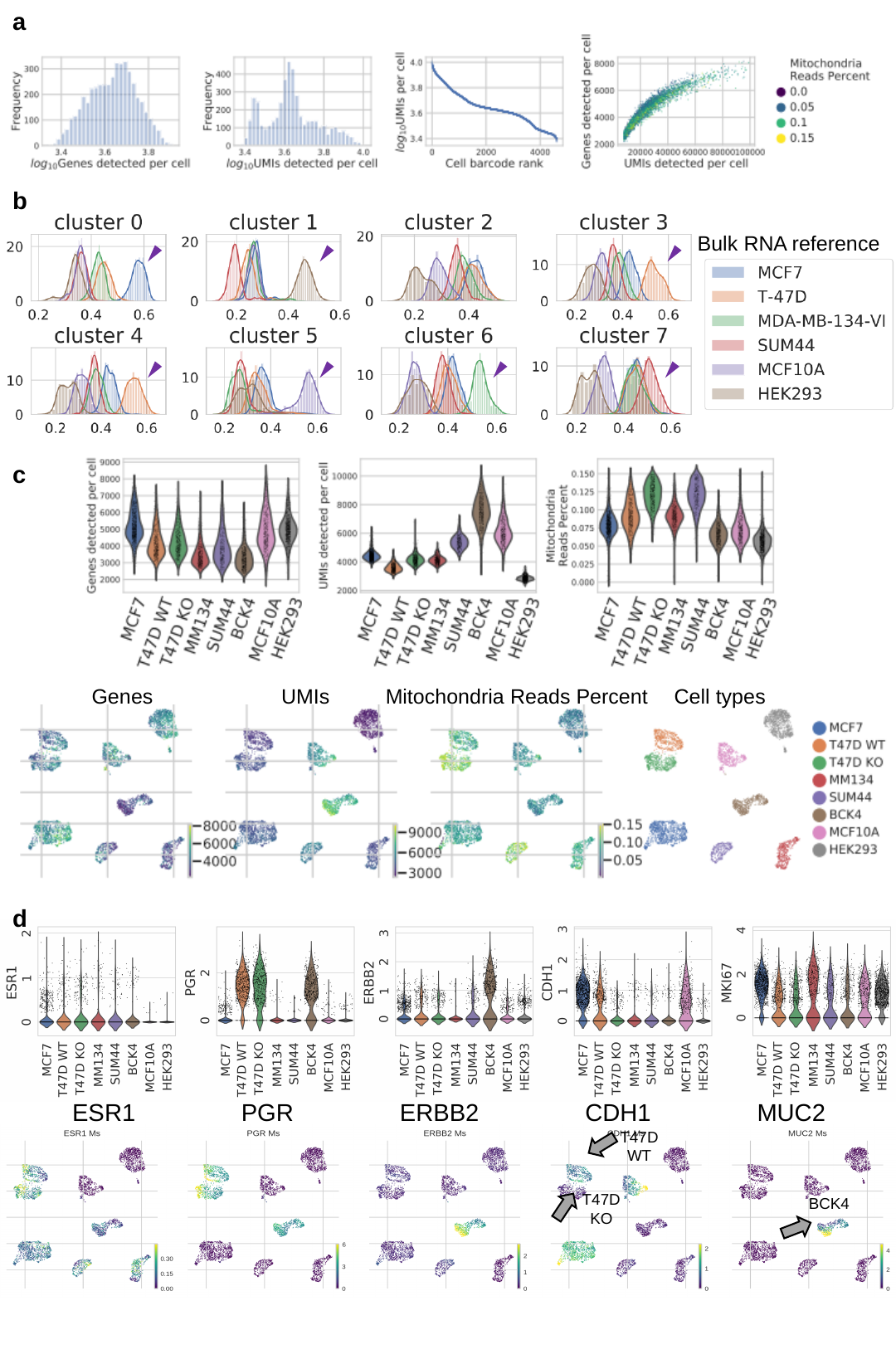

a
b
Bulk RNA reference
c
Genes
UMIs
Mitochondria Reads Percent
Cell types
d
ESR1
PGR
ERBB2
CDH1
MUC2
T47D WT
T47D KO
BCK4

### Slide 2
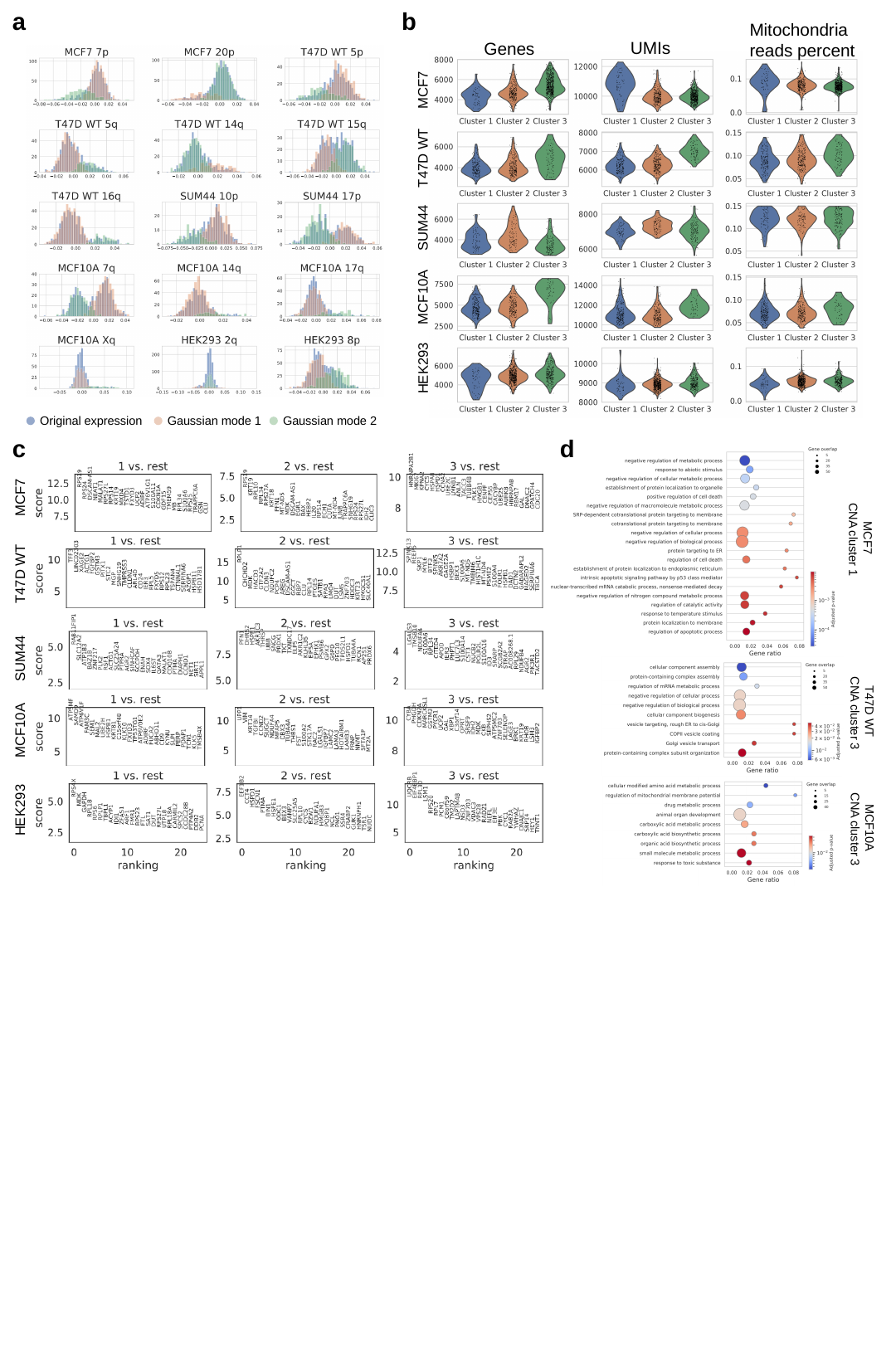

a
b
Mitochondria
reads percent
UMIs
Genes
MCF7
T47D WT
SUM44
MCF10A
HEK293
Gaussian mode 1
Gaussian mode 2
Original expression
d
c
MCF7
MCF7
 CNA cluster 1
T47D WT
SUM44
T47D WT
CNA cluster 3
MCF10A
HEK293
MCF10A
 CNA cluster 3

### Slide 3
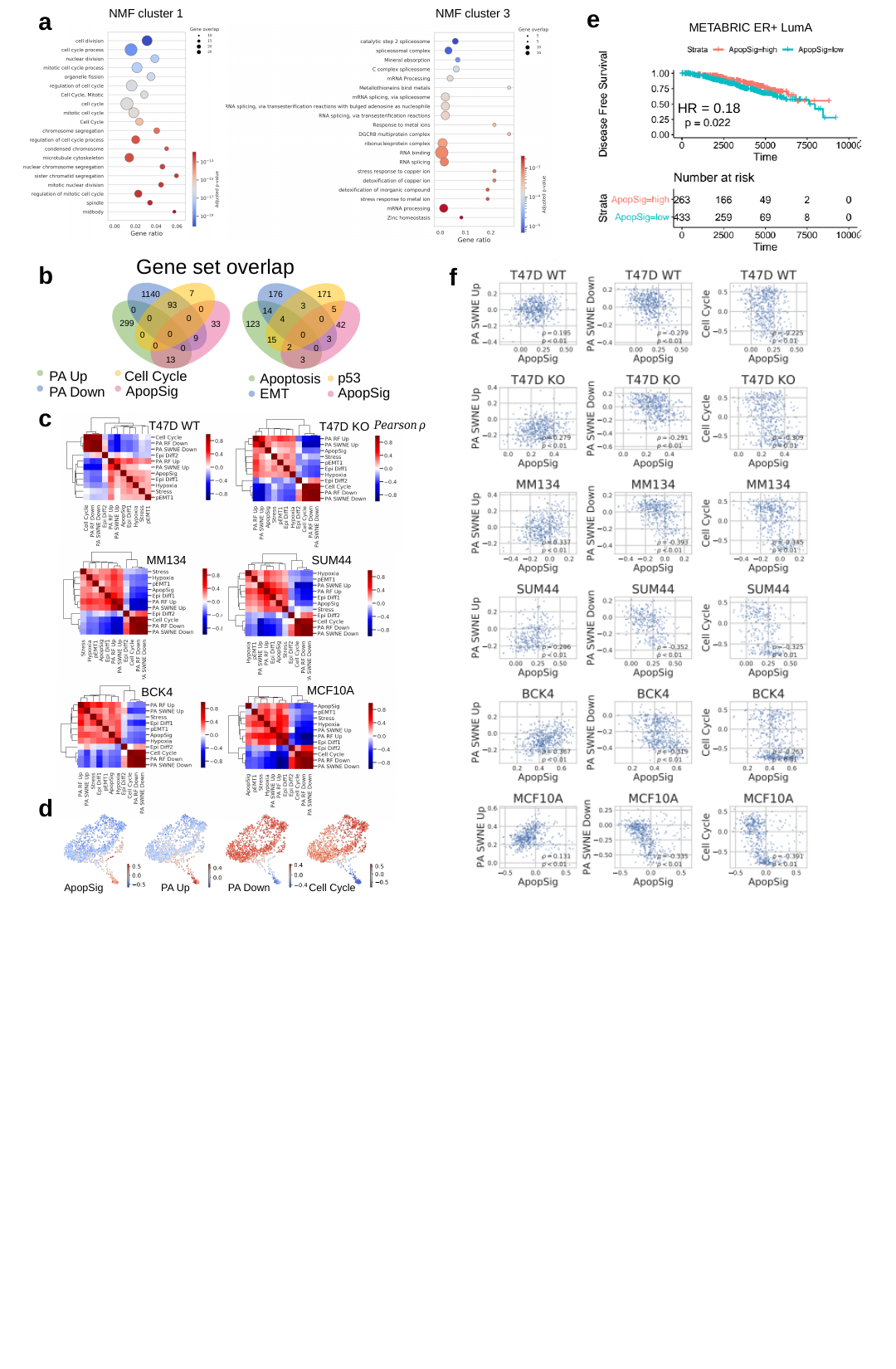

NMF cluster 1
NMF cluster 3
e
a
METABRIC ER+ LumA
HR = 0.18
HR = 0.28
Gene set overlap
b
f
7
1140
176
171
93
3
0
5
0
14
0
0
0
4
299
33
123
42
0
0
0
9
3
15
0
2
0
0
3
13
Cell Cycle
PA Up
p53
Apoptosis
ApopSig
PA Down
ApopSig
EMT
c
T47D WT
T47D KO
MM134
SUM44
MCF10A
BCK4
d
ApopSig
PA Up
PA Down
Cell Cycle

### Slide 4
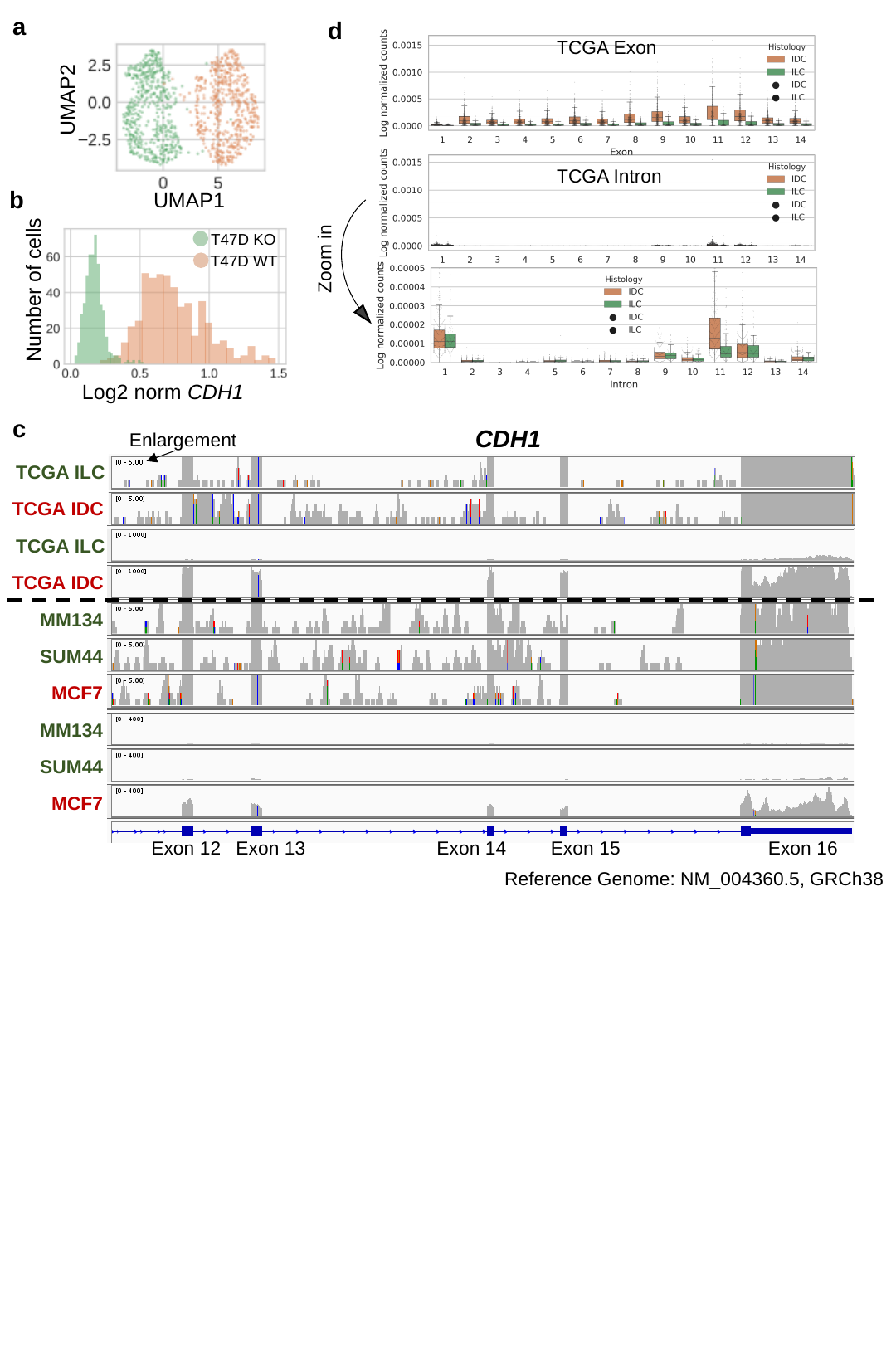

a
d
TCGA Exon
UMAP2
TCGA Intron
b
UMAP1
T47D KO
Zoom in
T47D WT
Number of cells
Log2 norm CDH1
c
CDH1
Enlargement
TCGA ILC
TCGA IDC
TCGA ILC
TCGA IDC
MM134
SUM44
MCF7
MM134
SUM44
MCF7
Exon 12
Exon 13
Exon 14
Exon 15
Exon 16
Reference Genome: NM_004360.5, GRCh38

### Slide 5
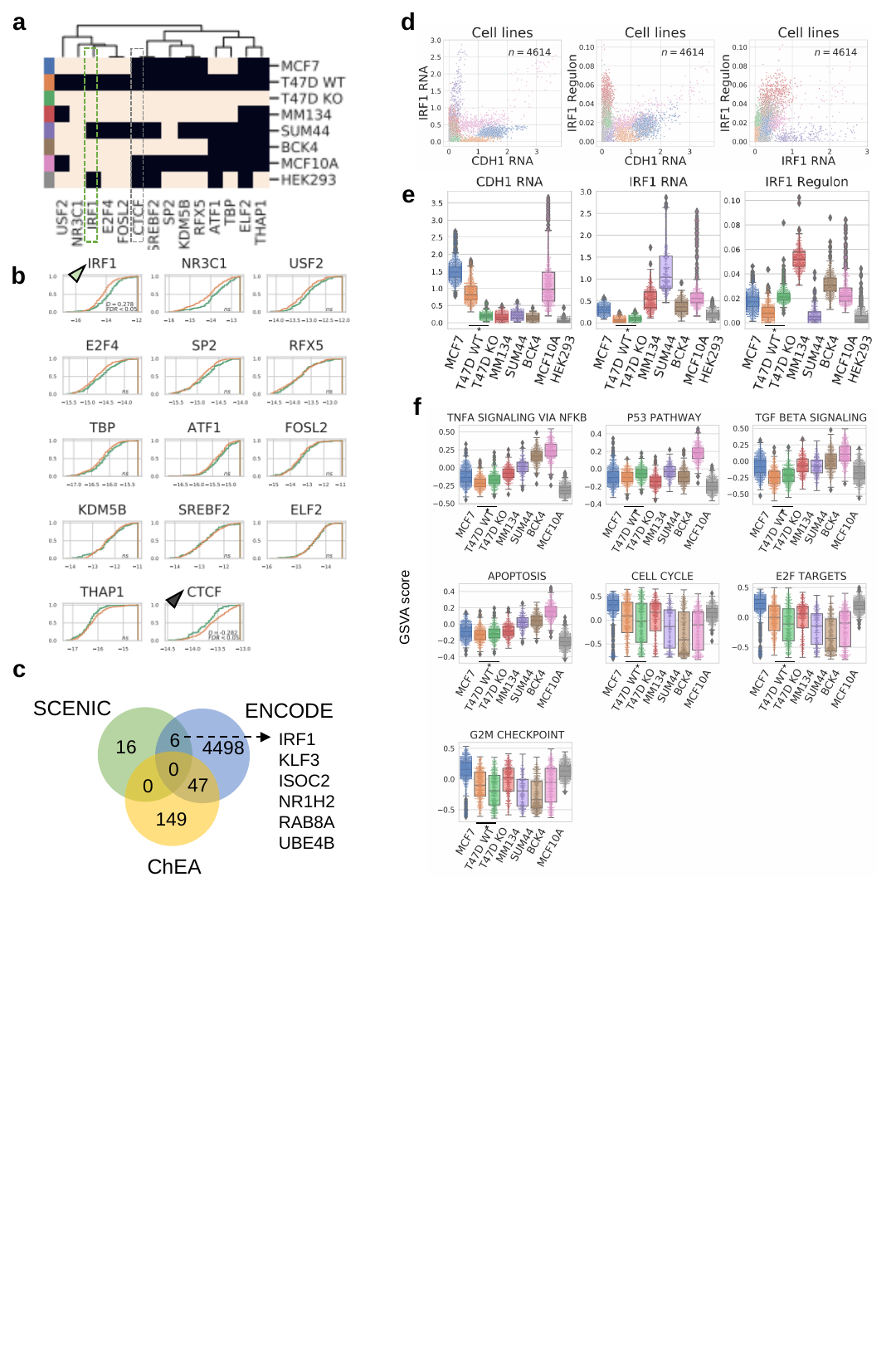

a
d
e
b
*
*
*
f
*
*
*
GSVA score
c
*
*
*
SCENIC
ENCODE
6
16
4498
0
47
0
149
ChEA
IRF1
KLF3
ISOC2
NR1H2
RAB8A
UBE4B
*
